# Histopathological Changes In The Intestine Of Infected Pigeons (*Columba Livia Domestica*) Caused By Helminthes Infection From Al-Qassim, Saudi Arabia

**DOI:** 10.1101/2021.12.18.473269

**Authors:** Mashael A. Aldamigh, Amaal H. Hassan, Ahlam A. Alahmadi

**Author notes:** **Funding** This research did not receive any specific grant from funding agencies in the public, commercial, or not-for-profit sectors. **Disclosure statement** The authors report there are no competing interests to declare.

## Abstract

Helminthes infection causes extensive harm to the pigeon host. The purpose of this study was to observe histopathological changes caused by helminths infection. Thirty-five pigeons (*C*.*L. domestica*) were purchased weekly from a bird’s market from Al-Qassim region, Saudi Arabia. Out of the 35 pigeons examined, 9 pigeons (25.71%) were found infected with helminth parasites, which were identified as one cestode (*Raillietina sp*.), and one nematode (*Ascaridia columbae*). The infected pigeons suffered from growth retardation, emaciation, weakness, droopiness, and diarrhea. A lot of histopathological changes were seen in the intestine of infected pigeons including atrophy and distortion of villi, infiltration of inflammatory lymphocytic cells, erosion, and loss of the typical structure of the intestine, necrosis in villi, and blood vessels congestion. This study concludes, for the first time in AL-Qassim region-Saudi Arabia, that the infection with helminth parasites caused significant histopathological changes in the intestines of the infected pigeons, and this could lead to increased mortality to the infected pigeons. Further work is necessary in Saudi Arabia to determine the prevalence and biological factors that have a significant impact on the helminth parasites community.

## Introduction

Pigeons are universal birds (Sari *et al*., 2008), and those of the order: *Columbi* sp. particularly (*Columba livia domestica*) are an important source of food (meat and eggs). Pigeons are hosts to a wide range of ecto- and endoparasites (Marques *et al*., 2007; Sivajothi and Reddy, 2014). Endoparasites consist of protozoan and helminth parasites (Ghazi *et al*., 2002). A suitable environment for the majority of helminthes in the intestine. The intestine provides them with food and safe shelter (Bernard and Matthews, 2001). In general, pigeons infected with helminth parasites can present several symptoms including emaciation, diarrhea, weakness, decreased growth, hemorrhage in the intestine, and digestive tract obstruction (Kaufmann, 1996). Additionally, infection with helminth parasites can cause considerable damage to the host’s tissue (Hoste, 2001), and may lead to death (Tanveer *et al*., 2011). Histopathology has a great importance in the medical diagnosis of diseases. A few studies reported histopathological changes in the infected pigeons with the helminth parasites. These histopathological changes include ulceration, enteritis, epithelial cell degeneration, destruction of epithelium secretory gland, and lymphocyte macrophage infiltration (Abed *et al*., 2014). In this study, the characterization of the histopathological changes in the intestines of pigeons after helminths infection will be presented.

## Materials and methods

Thirty-five pigeons (*C*.*L. domestica*) were purchased weekly from a bird’s market from AL-Qassim region, Saudi Arabia. Pigeons were dissected based on the procedure described by Al-Hussaini and Demian (1982). The intestines of pigeons were removed and examined for the presence of helminth parasites. For histopathological examination, tissue samples from the intestines of both infected and uninfected pigeons were collected and fixed in 10% buffered neutral formalin for 24 hours. After fixation, the tissue samples were washed with 70% alcohol, then gradually dehydrated in a serial concentration (70% - 100%) of alcohol. After dehydration, the tissue samples were cleared in xylene and then embedded in paraffin wax. Lastly, six-micron thick sections were prepared on slides for examination via routine histological techniques. The slides were placed on a hot plate with a temperature of 40°C to extend the sections, then the sections were stained by hematoxylin and eosin staining. The stained sections were identified and photographed using a light microscope according to the helminthological keys of Soulsby (1982) and Yamaguti (1961).

## Results

As a general observation, the infected pigeons suffered from growth retardation, emaciation, weakness, droopiness, and diarrhea (Figure 1A). Out of the 35 pigeons examined, 9 pigeons (25.71 %) were infected with helminth parasites (Figure 1B), which were identified to be one cestode species (*Raillietina* sp) and one nematode species (*Ascaridia columbae*) (Figure 1 E & F). No trematodes were recorded. Post dissection, the intestines of infected pigeons appeared to be thickened with mucus excretion because of a great number of helminths as compared to the healthy intestines. The hemorrhagic hematoma was observed in several locations throughout the gut of the infected intestines (Figure 1 C & D). Histologically, the normal structure of the bird’s intestine is multi-layered composed of: mucosal layer, submucosal layer, circular muscle, longitudinal muscle, and serosal layer. The anterior surface of the intestine with a complexed and folded structure known as villi, which increases the surface range and absorbent area of the intestinal tract (Figure 2 A). Several histopathological changes associated with helminths infection were observed. These changes included atrophy and distortion of villi and glands, infiltration of inflammatory lymphocytic cells, desquamation of the lining epithelium inside the lumen with erosion, and loss of the typical structure of the intestine (Figure 2 B). Additionally, blood vessels showed congestion and mild thickening of the vessel’s walls (Figure 2 C). The intestines revealed necrosis of the intestinal mucosa at different points (Figure 2D).

**Figure 1.**
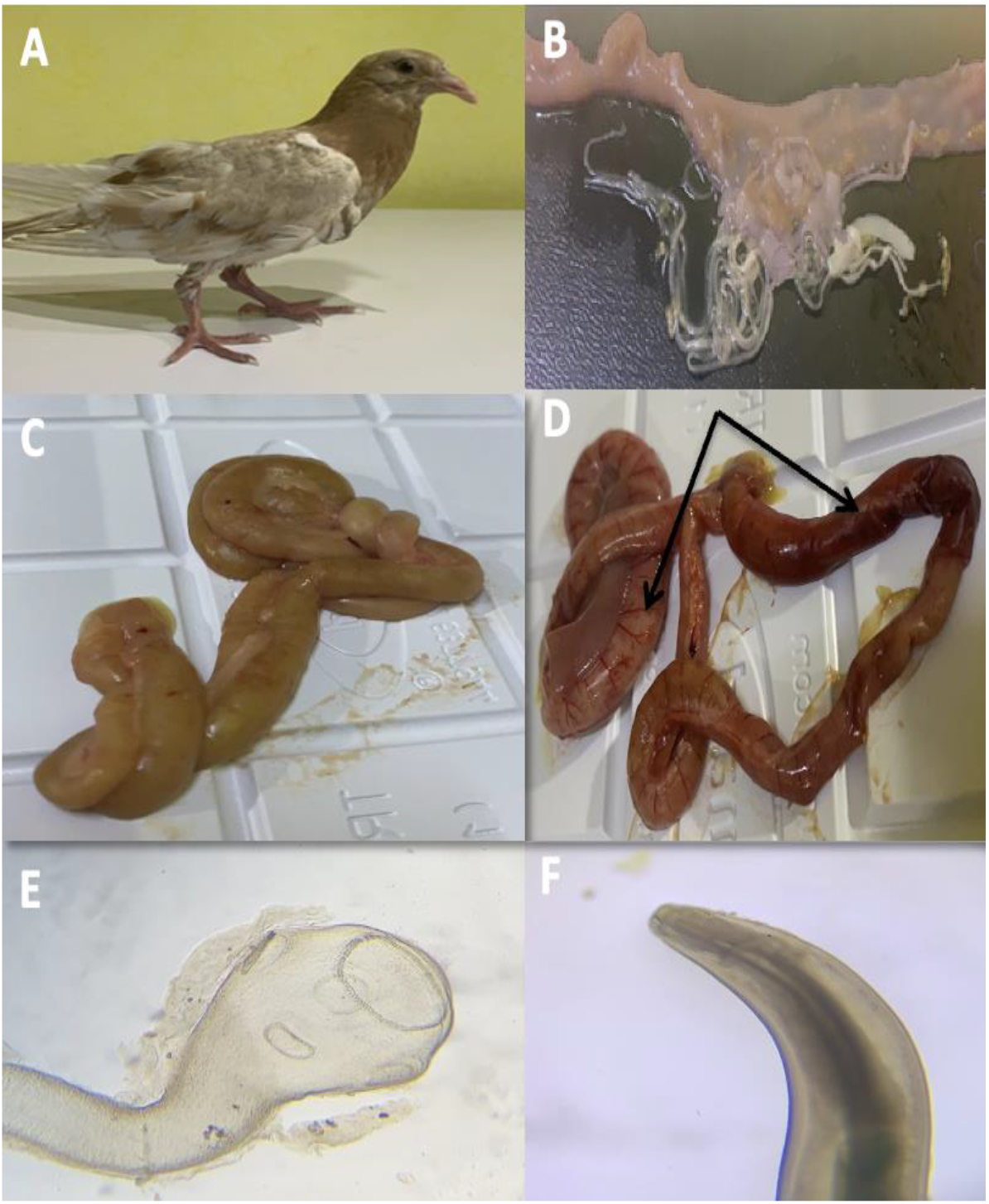
A. Infected pigeon suffered from emaciation, diarrhea, and growth retardation. B. The intestine of infected pigeon showing infection of helminth parasites. C. The intestine of uninfected pigeon. D. The intestine of infected pigeon showing haemorrhagic haematoma in several sections. E. Cestode (*Raillietina* sp.) under light microscope (10X). F. Nematode (*Ascaridia columbae*) under light microscope (10X)

**Figure 2.**
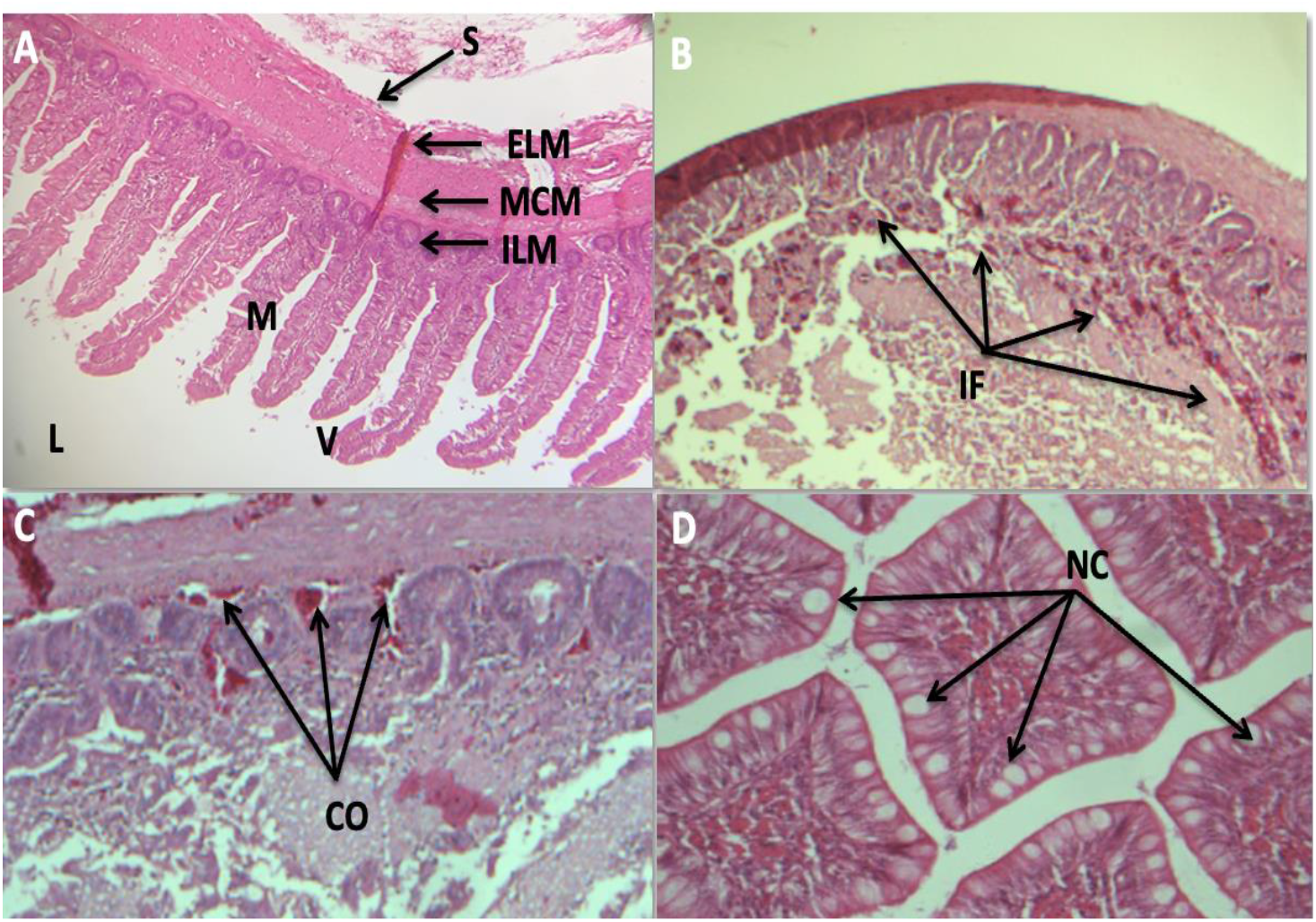
A. Cross section in the intestine of uninfected pigeon revealing different intestine layers: Serosa (S), muscularis (LML: an outer longitudinal muscle layer, CML: a middle circular muscle layer and LML: an inner longitudinal muscle layer), mucosa (M), villi (V) and lumen (L) (X10) H & E stain. B. Cross section in the intestine of infected pigeon showing distortion of villi with erosion and infiltration of inflammatory lymphocytic cells (IF) (X10) H & E stain. C. Cross section of infected intestine showing blood vessels congestion (CO) (20X) H & E stain. D.Cross section in the intestine of infected pigeon showing necrotic tissue (NC) in some damaged parts of mucosa (40X) H & E stain.

## Discussion

Histopathological changes in the intestine can be used as a diagnostic tool for infected pigeons with helminth parasites. The objective of this study was to demonstrate the histopathological changes in the intestine of pigeons after helminthic infection for the first time in AL-Qassim region, Saudi Arabia. In accordance with previously reported findings (Kamal *et al*., 2020), the intestines of the infected pigeons were completely blocked by a great number of helminth parasites. This blockage led to the development of hemorrhagic hematoma at different points in the intestinal tract. Moreover, the blood vessels were congested with mild thickening of their walls. The current study confirms that helminths infection has an obvious harmful effect on the host pigeon. The infected pigeons suffered from growth retardation, emaciation, weakness, and diarrhea, as previously reported by (Kaufmann,1996). The most prominent histopathological changes in the intestinal tissue of the infected pigeons were the abrasion and morphological changes in the whole structural design of the intestine. These changes include atrophy and distortion of villi and glands with erosion, losing the typical structure of the intestine, infiltration of inflammatory lymphocytic cells, and increased necrosis of the intestinal mucosa with desquamation of the lining epithelium inside the lumen. These findings were supported by the previously reported histopathological changes in the intestines of pigeons infected with *R*.*tetragona Molin* 1858 (Cestode) and *A*.*columbae Gmelin*, 1780 (Nematode) (Shaikh *et al*., 2016). In this study, the authors observed the architectural disintegration of the muscular layer, destruction of Brunner’s and crypt glands, necrosis of serosal layer, migratory tunnels formed along with fibrosis, atrophy of villi, and infiltration of mononuclear (macrophages and lymphocytes) in lamina propria. Moreover, these histopathological changes are in accordance with the previously reported findings in the intestines of domestic pigeons (*C*.*L. domestica*) caused by helminths infection in Egypt (Ibrahim *et al*., 2018), and the intestines of domestic chickens that infected with the cestode *Cotugnia* sp. from Iraq (Mahdi *et at*.,2018).

## Conclusion

This study concludes, for the first time in AL-Qassim region-Saudi Arabia, that the infection with helminth parasites caused significant histopathological changes in the intestines of the infected pigeons. Consequently, this could lead to increased mortality to the infected pigeons. Further work is necessary in Saudi Arabia to determine the prevalence and biological factors that have a significant impact on the helminth parasites community.

## Acknowledgment

The authors are grateful to the Department of biology at Faculty of Science at King

Abdulaziz University, for providing the facilities to carry out this study.

